# Thermoregulation and Diurnal Roost Selection of Boreal Bats During Pre-Hibernation Period

**DOI:** 10.1101/2024.05.22.595441

**Authors:** Kati M. Suominen, Niclas R. Fritzén, Mari A. Fjelldal, Anna S. Blomberg, Minna J.K. Viljamaa, Thomas M. Lilley

## Abstract

Living in a seasonal environment poses challenges for small mammals, such as bats, reliant on insects as their primary food source. Bats may adeptly navigate these energetic challenges by reducing their metabolism and body temperature, entering a state of torpor. Particularly during the winter, bats remain torpid for extended periods, but are dependent on sufficient energy reserves to survive until spring. With the onset of autumn and declining temperature, bats face the challenge of building their fat deposits during a time of decreasing food availability. Bats may therefore transition to cooler roosts to initiate torpor, thereby reducing energy expenditure. However, little is still known about torpor use or roost selection by bats in autumn. This study explores the factors influencing roost selection and torpor use and –duration in two bat species during this critical transition period between the breeding and overwintering season. We show that date in autumn is a stronger driver of torpor use than prevailing ambient temperature, and that bats employ specific strategies in which they first increase daytime torpor use before also increasing the use of night-time torpor during the pre-hibernation fattening period, most likely to facilitate rapid fat accumulation. Notably, bats commenced night-time torpor use after spending entire days in torpor. These findings underscore the dynamic nature of torpor and the energy-saving strategies employed during the crucial pre-hibernation period, marking the transition from summer to winter.

## Introduction

Organisms inhabiting high latitudes confront challenges associated with pronounced seasonality, featuring long and cold winters with limited food resources, contrasted by brief summer seasons. These provide a constrained timeframe for reproductive activities and preparations for the ensuing prolonged and harsh winter. The summer period is short with pronounced ambient light conditions, thereby presenting additional challenges for nocturnal animals, particularly those dependent on swarming insects, such as bats (Chiroptera). As nights become longer with the approaching autumn, bats engage in energy accumulation to endure the forthcoming winter; however, this period coincides with a decline in temperature, which in turn is associated with insect biomass decline (Welti et al., 2021), causing an energetic challenge for bats. Nevertheless, bats, along with certain other small vertebrates, have surmounted these challenges through the strategic utilization of *torpor* as an adaptive mechanism to contend with unfavourable environmental conditions (Speakman & Thomas, 2003; Geiser, 1988, Geiser, 2004; Geiser, 2013).

Torpor, defined as a state of diminished physiological activity in an organism, is characterized by a reduction in metabolic rate, body temperature, and overall energy expenditure (Barclay et al., 2001). This transient condition of dormancy or inactivity serves to conserve energy (Speakman & Rowland, 1999) or water (Bondarenco et al., 2016; Muñoz-Garcia et al., 2022) when resources are scarce or environmental conditions are adverse. Importantly, torpor is a reversible state, allowing swift return to the normal active state when conditions ameliorate (Geiser & Ruf, 1995). The use of torpor demonstrates seasonal and diurnal variations. During winter, bats implement extended torpor bouts and reduced metabolism, allowing for the conservation of energy reserves until the resurgence of insects in spring (Speakman & Thomas, 2003; Czenze et al., 2017; Boyles et al., 2020). In summer, the use of torpor is often associated with environmental factors such as ambient temperature, (Geiser & Bringham, 2000, Fjelldal et al., 2021; Fjelldal et al., 2023a), precipitation, barometric pressure and overall climatical conditions (Ruf & Geiser, 1995), either by directly affecting the energy reserves (Speakman & Rowland, 1999) or through reduced food availability (Poulsen, 1996; Speakman et al., 2000; Ruf & Geiser 2015; Welti et al., 2022).

Torpor patterns can vary among bat species, influenced by factors such as their ecological niche, geographic location, and the availability of resources (Stawski & Geiser, 2010; Boyles et al., 2017). Bats with a greater body mass exit torpor at lower ambient temperatures, whereas those with a lower body mass delay arousal until temperatures are higher (Sørås et al., 2022). This implies that bats may, based on their individual state, adjust their level of thermoregulation at different temperatures, potentially to alleviate the associated costs of torpor (Sørås et al., 2022) or to obtain physiological or ecological benefits from being active (Fjelldal et al., 2023b).

Changes in torpor use are also reflected in the roost site selection (Speakman & Rowland, 1999), and it has been suggested that thermal physiology might have co-evolved with roost preference (Czenze et al., 2022). In the summer, during breeding time, bats choose warmer roosts (Speakman & Rowland, 1999) and when ambient temperature within the shelter is high, the physiological and ecological costs of torpor may exceed the benefits, hence the animal should maintain higher body temperature and metabolism (Boyles et al., 2020). However, at lower temperatures within roosting environments, bats undergoing torpor can conserve approximately one-third of energy compared to those roosting in warmer conditions, thereby establishing the advantageous nature of roosting at colder temperatures and utilizing torpor during the pre-hibernation season (Speakman & Rowland, 1999). Intensified torpor use, in combination with increased foraging rates and reduced metabolic signalling of leptin during the pre-hibernation fattening period are all mechanisms believed to contribute to the rapid fat accumulation observed in many bat species (Kunz et al., 1998; Speakman et al., 1999; Kronfeld-Shcor et al., 2000; Fraser & McGuire, 2023).

Nevertheless, the specific cue prompting bats to transition to colder roosts to engage in torpor remains unknown. It is plausible that air temperature serves as a cue, signalling a scarcity in food resources for insectivorous bats, particularly when ambient temperatures fall below 14°C (Speakmann et al., 2000). Changes to the photoperiod have also been observed to result in individuals optimizing energy conservation through torpor and energy intake via food consumption, contributing to enhanced survival in marsupials (Turner & Geiser, 2016). These changes to photoperiod, evidenced as increasing night length with date since summer solstice at northerly latitudes, may also influence changes in torpor expressions by bats in autumn; however, the use of torpor by bats throughout the pre-hibernation fattening period is largely understudied.

The objective of our study was to investigate thermoregulatory strategies in the two boreal bat species, *Eptesicus nilssonii* and *Myotis brandtii*, with differing ecology, during the pre-hibernation period. We aimed to investigate whether ambient temperature (T_a_) or date in autumn serves as the more influential predictor of daily and nightly torpor use in the critical transition period from summer to winter. While we acknowledge that insect abundance, and therefore bat activity, is influenced by temperature and weather conditions (Speakman et al., 2000; Welti et al., 2022), we hypothesize that date would exert a stronger influence on torpor expressions in boreal bats. This is based on the expectation that the gradual decrease in available insects in autumn is not solely driven by decreasing T_a_ but also by phenology. Additionally, as bats accumulate more fat reserves throughout autumn, state-dependent torpor responses can alter the influence of weather conditions on individuals. In this study we also describe the diurnal roost types used by these species in autumn and hypothesize that roosts within built structures and trees offer higher T_a_, but less stability compared to those in rock screes, consequently leading to lower skin temperatures (T_sk_) during torpor in boulder fields. We further posit the presence of species-specific differentials in both torpor strategies and roost selection, arising from inherent variances in species biology. *E. nilssonii* hibernates in colder conditions compared to *M. brandtii* (Masing & Lutsar, 2007), hence we expect it to also utilise colder roosts during the pre-hibernation period.

## Material and methods

### Study species

Our study species, *E. nilssonii* and *M. brandtii*, are widely spread across Northern Europe. The distribution range of *E. nilssonii* ranges from Europe to Asia and Japan (Suominen et al. 2022). Of all bat species, it has the northernmost distribution area in the world and is often observed above the Arctic Circle (Tidenberg et al., 2019, Kotila et al., 2022). *Myotis brandtii* occurs in Central and Northern Europe with the distribution range extending to the Ural Mountains (IUCN). Both species are insectivorous. *Eptesicus nilssonii* is slightly larger than *M. brandtii*, weighing between 10 to 20 grams, whereas *M. brandtii* typically weighs 6 to 10 grams.

### Research area

Our research was conducted on Valsörarna islands, with the main islands of an approximately 4 km_2_ situated at 63°25’N 21° 5′E in the outer archipelago of Kvarken, Baltic Sea, between Finland and Sweden. Despite its relatively modest size, the islands boast a diverse landscape. Extensive areas feature open heaths with exposed boulder fields, while the forested regions consist mainly of deciduous forests dominated by birch and aspen with alder and rowan along the shores. Evergreen trees are limited to small groups or individual specimens. Due to minimal human influence over an extended period, the area is characterized by a substantial presence of dead and decaying trees. The landscape also includes various forested and open mires, shore meadows, and coastal lagoons. The absence of a permanent settlement is notable, with only a few predominantly old wooden cottages present. The primary islands, along with its surrounding islets, is part of a privately owned nature reserve, forming a crucial route for migrating birds and bats.

### Bat capture and handling

Bats were captured from August 9^th^ to September 18^th^, and T_sk_ and location data were collected between August 9^th^ and October 1^st^, 2021. We captured bats using harp traps, which were checked every 30 minutes. Additionally, there were seven accessible bat boxes in the area, from which we collected potential roosting bats daily. For each bat, we performed species identification and determined their sex, weight, forearm length, age class (Anthony, 1988), and reproductive state (Mitchell-Jones & McLeish 2003). Subsequently, we attached a radio transmitter with a temperature sensor (Telemetrie-Service Dessau, Telemetriesender V4 Temperatur) to each bat using skin glue (Sauer-Hautkleber). The temperature-sensing transmitters weighed 0.55 g with a minimum lifespan of 42 days for *E. nilssonii*, and 0.45 g with a minimum lifespan of 21 days for *M. brandtii*. The tag weight was maintained below 5% of the bat’s body mass, adhering to the commonly accepted maximum carrying load for bats (Aldridge & Brigham, 1988). These tags adjust time intervals between pulses based on T_sk_, providing information on the T_sk_ of the bats. Following the application of the transmitter, we placed each bat in cotton cloth for 15 minutes to allow the adhesive to dry, after which we released the bats at the capture site, following the protocol established by Mitchell-Jones & McLeish (2003).

### Radio receiver network system

An automatic radio-tracking system of ten RTEU-radio stations was established on the island in the summers of 2020 and 2021. Each station consists of a 6-meter-tall pole equipped with four directional antennas (either Moxon or H-antennas), each covering a 90° sector. The antennas are connected to a RaspberryPi computer through an individual USB-radioreceiver (Nooelec NESDR). To enable remote access, each station was equipped with an internet connection. Further details about the RTEU-system can be found in Gottwald et al. (2019). Power for the system was derived from solar panels (140 or 150 W) and a 100 Ah AGM battery.

The direction from a radio station to the transmitter was determined by assessing the relative gains of two neighboring antennas. The tag’s position was approximated by the point of intersection of two lines obtained from bearing calculations at two distinct stations. In cases where more than two stations concurrently received the signal, the centroid of the resulting polygon was calculated (refer to Gottwald et al., 2019, for additional details). Individual tags were differentiated based on their specific frequencies. Every morning, data from the preceding night were downloaded from all stations, processed, and analyzed to identify signals corresponding to each previously used tag. Maps illustrating the intersection points or polygon centroids were then generated. Using early morning data, we approximated the area each bat had traversed. Subsequently, we conducted manual searches in the field using handheld radio receivers (Televilt RX98 and ICOM IC-R30) to locate the day roosts of each bat.

### Data management

T_sk_ data were converted with 10-minute intervals from the recorded pulse frequencies, and we applied the methods described in Fjelldal et al. (2023a) to identify torpor bouts and phases (i.e. ‘torpor entries’, ‘stable torpor’ and ‘torpor arousals’). To apply this method, we initially calculated species-specific torpor thresholds using the equation for determining torpor onset values in small mammals by Willis (2007). The calculated torpor onset values were 30.0°C for *E. nilssonii* and 29.9°C for *M. brandtii* after subtracting 2°C (due to the difference between body temperature (T_b_) and T_sk_ generally being ∼2°C; Audet & Thomas, 1996). After applying the methods for phase determination, we performed the suggested sensitivity analyses (Fjelldal et al., 2023a) to adjust ΔT_sk_ threshold values accordingly. When considering torpor in the analyses we excluded arousal phases as these are energetically more resembling the euthermic state than torpor (Geiser et al., 2014).

Weather data were downloaded with 10-minute intervals from the meteorological stations Korsholm, Valsörarna (T_a_, wind speed and barometric pressure) and Vasa Klemetsö (precipitation) through Finnish meteorological institute open data source (FMI). Sunrise and sunset times were obtained through the package suncalc (Thieurmel & Elmarhraoui, 2019) in R.

### Statistics

#### Temporal trends in torpor use

We performed all analyses in R (version 4.3.1). To determine whether T_a_ or day in autumn (days since 1^st^ August) were the main driver of torpor use throughout autumn in the two species, we calculated the proportion spent torpid for each day (from sunrise to sunset) and night (from sunset to sunrise). However, initial visualisations of the data indicated an apparent shift in torpor use by the end of August that did not seem to correspond to measured T_a_ (Fig. S1 in Supplementary Materials 1). Because we were interested in potential changes in torpor use strategies between summer and the pre-hibernation period, we performed a Davies’ test (package segmented; Muggeo 2008) on a simple binomial generalized linear model with the proportion spent torpid as the response and day in autumn and time of day (day/night) as predictors, to identify the presence of a change in the slope. The Davies’ test was significant and indicated a breakpoint around 29 days after 1^st^ August. To obtain the exact breakpoint estimate, we then performed a breakpoint analysis (package segmented) with an initial starting value of 29, which resulted in a final breakpoint estimate of 28 days after 1^st^ August (Fig. S2 in Supplementary Materials 1). Because of this apparent shift in temporal torpor use we chose to analyse the period prior to and after this point in time separately (referred to as ‘late summer period’ and ‘autumn period’ onwards).

For each period we fitted binomial generalized mixed models (package lme4; Bates et al., 2015) with the proportion spent torpid as response variable, and either mean, min or max T_a_ or day in autumn (these could not be tested together due to high correlations) in addition to species, time of day (night/day), roost type, sex (this could not be included for the late summer period as only one observation was from a female), total rainfall, mean windspeed, and mean barometric pressure as predictors. Due to few datapoints in the late summer period (N_obs_ = 43), only time of day and T_a_ or date was tested as interaction effects. For the autumn period we tested interactions between each weather (or date) variable and species, time of day, roost type and sex (N_obs_ = 228). Individual ID was included as a random effect in all models. We then performed a model selection on the global models using the *dredge* function from the MuMIn package (Barton & Barton, 2015). The AICc of the highest ranked model for each T_a_ variable and date variable were then compared to determine the best fit to the data for each of the two periods.

#### Roost type effects on torpor dynamics

To test whether the type of day roost impacted torpor dynamics in *E. nilssonii* and *M. brandtii*, we constructed mixed models with either torpor bout durations (hours; glmer with Gamma distribution and log link; N_obs_ = 210) or minimum T_sk_ (lmer; N_obs_ = 137; only testing torpor bouts that were > 1 hour in duration to allow T_sk_ time to decrease) during each bout as response variables. Roost type, species, sex, minimum T_a_, mean wind speed and mean barometric pressure were included as predictors, and we tested roost type, species, and sex in interaction with each weather variable in addition to interactions between roost type, species, and sex. ID was included as a random effect. We performed model selections on the global models as described in the previous paragraph.

## Results

### Captured bats and day roosts

In total, we captured 14 *E. nilssonii* and 24 *M. brandtii* (Table 1). Of these individuals, after being radio tagged, three *E. nilssonii* and eight *M. brandtii* could not be relocated (they either left the research area or dropped the tag out of reach of the radio stations), and therefore this study includes data on 11 *E. nilssonii* and 16 *M. brandtii*. In total we recorded roost choices for 165 days for *E. nilssonii* and 70 for *M. brandtii*. The bats utilized three main diurnal roost types: buildings, tree cavities and boulder fields (Table 1).

**Table 1:**
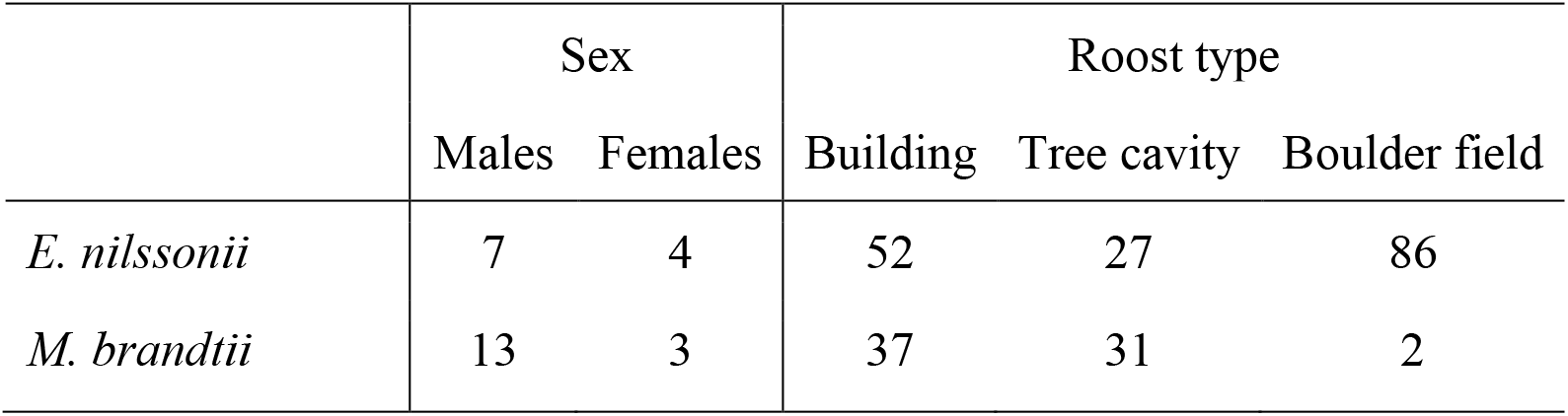
Diurnal day roost types utilized by *E. nilssonii* and *M. brandtii*. The numbers under the roost types are number of days spent in each roost type per species.

|  | Sex |  | Roost type |  |  |
| --- | --- | --- | --- | --- | --- |
|  | Males | Females | Building | Tree cavity | Boulder field |
| *E. nilssonii* | 7 | 4 | 52 | 27 | 86 |
| *M. brandtii* | 13 | 3 | 37 | 31 | 2 |

To assess whether the selection of day roosts changed throughout autumn, we calculated a 10-day moving average of the proportion of roost choices for each day. Proportions based on fewer than 5 observations the last ten days were excluded. The results indicated that tree-roosts for both species were more frequently used in the early autumn but were exchanged for buildings and/or boulder fields as autumn progressed (Fig. 1).

**Figure 1:**
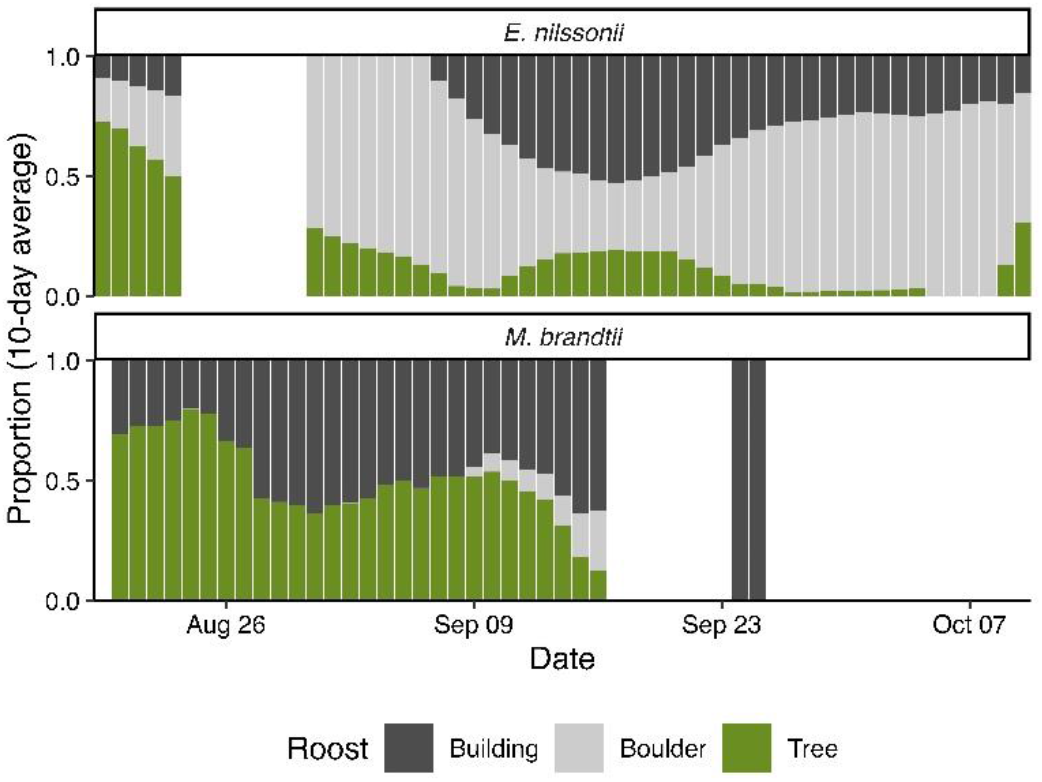
Proportions of roost types selected throughout the study period, based on 10-day moving averages.

### Temporal trends in torpor use

The highest ranked late summer model for explaining torpor use (proportion) only included time of day as predictor (predicted proportion of daytime torpor use: 0.83 [0.62-0.93 CI]; predicted proportion of night-time torpor use: 0.05 [0.01-0.27 CI]) (see Supplementary Materials 2 for model selection). Neither T_a_ nor date thus explained any of the variation in torpor use observed in late summer (from 09^th^ to 28^th^ of August).

For the autumn model (from 29^th^ of August to 09^th^ of October), however, the highest ranked model included date, time of day, rain, and wind as predictors (Table 2). Date was a drastically better predictor than any of the three T_a_ variables (AICc of highest ranked date-model = 88.7 versus AICc of highest ranked T_a_ -model = 111.2). Bats in autumn, independent of species and sex, used more torpor during daytime than at night and increased torpor use in response to later dates, higher wind speed and more rainfall. In response to date, bats appeared to first increase torpor use in the daytime from the end of August and generally spent the full day torpid already by early September (Fig. 2). However, they did not begin to increase nightly torpor use until they were already spending entire days in torpor.

**Table 2:** Predicted effects from the highest ranked logistic model explaining daily and nightly torpor use proportion in the autumn period. *Note*: The effect of night-time is the difference in effect size from the daytime effect (intercept).

| Variable | Random effects<br>Variance (SD) | Fixed effects |  |
| --- | --- | --- | --- |
|  |  | Estimate (SE) | *p* value |
| *ID* | 4.5×10<sup>-9</sup> (6.7×10<sup>-5</sup>) |  |  |
| Intercept (daytime) |  | -18.90 (± 3.28) | < 0.001 |
| Nighttime |  | -5.75 (± 1.03) | < 0.001 |
| Days since August 1<sup>st</sup> |  | 0.54 (± 0.09) | < 0.001 |
| Mean wind (m/s) |  | 0.32 (± 0.11) | < 0.01 |
| Total rainfall (mm) |  | 0.26 (± 0.09) | < 0.01 |

**Figure 2:**
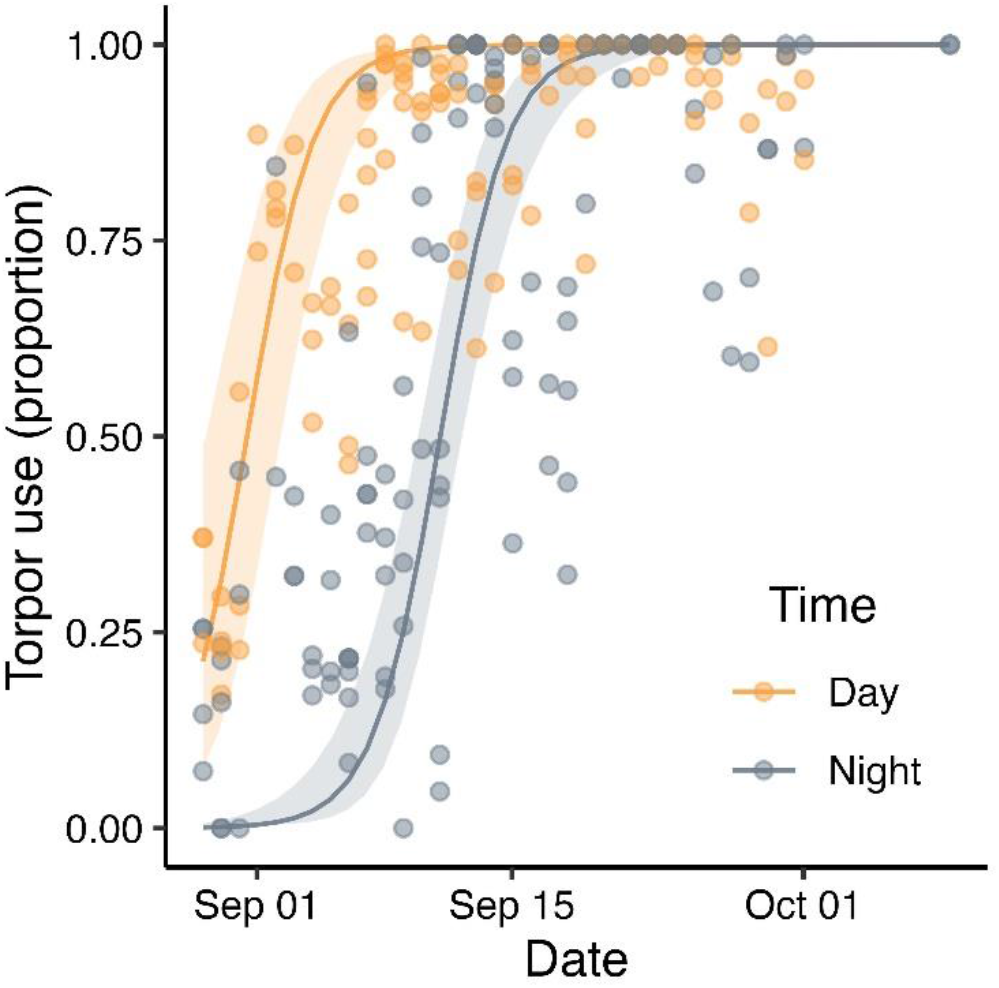
Proportion of torpor use as a function of date in autumn during day (yellow) and night (blue). The logistic regression lines are the predicted effects from the best model and thus account for rain- and wind-effects, while datapoints show the raw data.

### Roost type effects on torpor dynamics

We recorded a total of 213 complete torpor bouts, lasting from 20 minutes to 6.8 days. However, only three torpor bouts lasted longer than 51 hours (lasting 95.8 hours, 100.2 hours, and 161.5 hours, respectively) and were therefore excluded from the analyses. The mean torpor bout duration (disregarding the three longest bouts) was 8.5 hours (±10.5 *SD*). The highest ranked model (see Table S3 in Supplementary Materials 2 for model selection) included species, minimum T_a_ and barometric pressure as predictors. Roost type therefore did not significantly influence torpor bout duration; however, *M. brandtii* was predicted to employ generally shorter torpor bouts than *E. nilssonii*, and decreasing barometric pressure and T_a_ led bats to extend their torpor bouts (Table 3a & Fig. 3a).

**Table 3:** Predicted effects from the highest ranked models explaining **a)** torpor bout duration and **b)** minimum T_sk_ during torpor bouts. *Note*: The effect of species and roost types are the differences from the intercept.

| Variable | Random effects<br>Variance (SD) | Fixed effects |  |
| --- | --- | --- | --- |
|  |  | Estimate (SE) | *p* value |
| **a) Model: Torpor bout duration (hours)** |  |  |  |
| ID | 0.06 (0.24) |  |  |
| Residual | 0.85 (0.92) |  |  |
| Intercept (*E. nilssonii*) | | 29.73 ($\pm 2.36$) | < 0.001 |
| *M. brandtii* | | -0.61 ($\pm 0.23$) | < 0.01 |
| Minimum $T_a$ | | -0.21 ($\pm 0.03$) | < 0.001 |
| Mean hPa | | -0.025 ($\pm 0.002$) | < 0.001 |
| **b) Model: Minimum $T_{sk}$ during torpor bout** |  |  |  |
| ID | 0.00 (0.00) |  |  |
| Residual | 1049 (3.24) |  |  |
| Intercept (*E. nilssonii* \| building-roost) | | 5.56 ($\pm 0.07$) | < 0.001 |
| *M. brandtii* | | -1.30 ($\pm 0.74$) | 0.09 |
| Tree-roost | | -1.68 ($\pm 0.04$) | < 0.05 |
| Rock-roost | | -2.35 ($\pm 0.05$) | < 0.01 |
| Minimum $T_a$ | | 0.96 ($\pm 0.09$) | < 0.001 |

**Figure 3:**
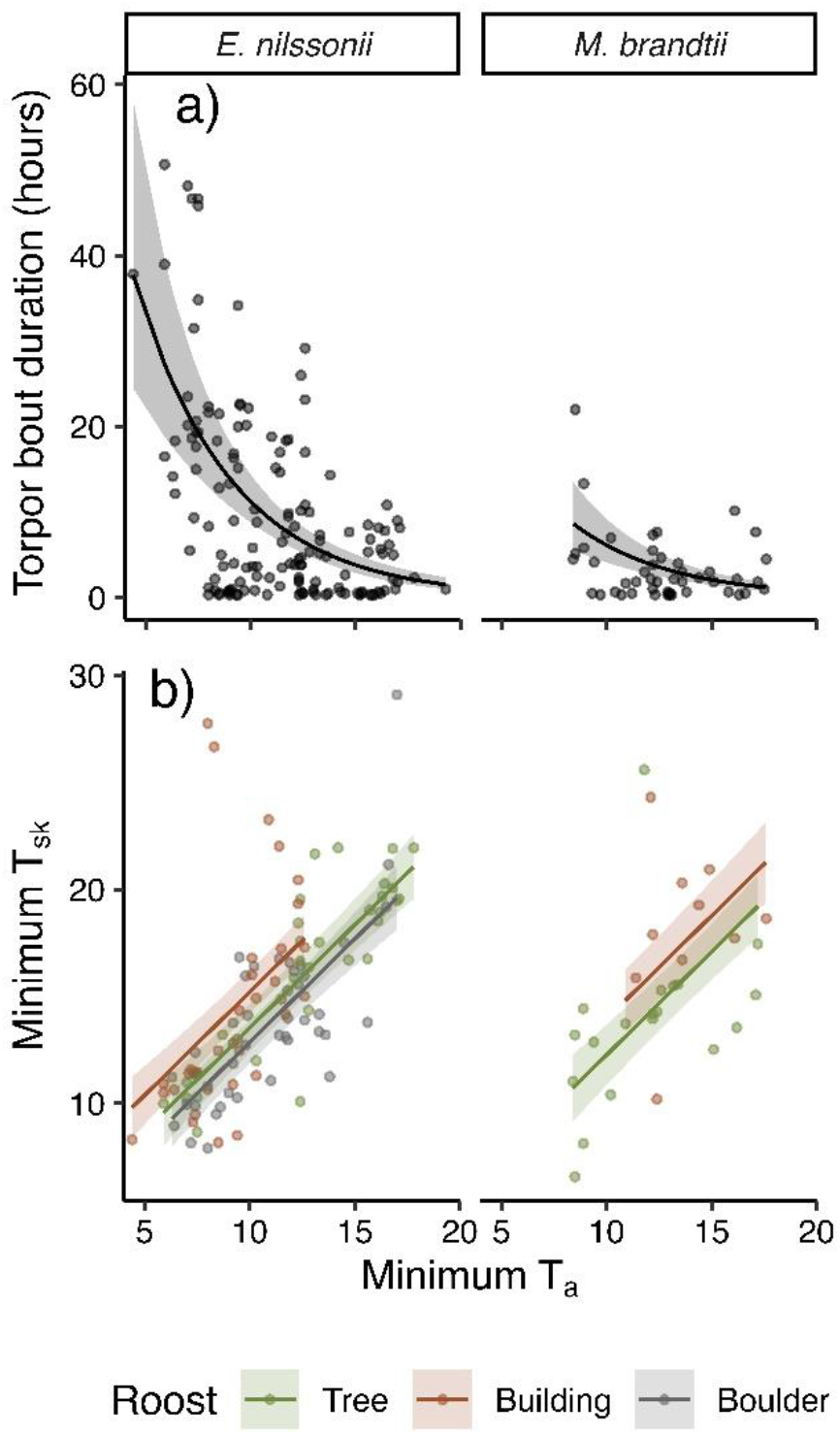
The predicted effects of minimum T_a_ on torpor bout dynamics. **a)** T_a_ effects on torpor bout durations for each species. **b)** The predicted effects of minimum a T_a_ on minimum T_sk_ during torpor for each species and roost types. Regression lines are the predicted effects from the best models while datapoints show the raw data.

The highest ranked model explaining minimum T_sk_ during torpor included roost type, species, and minimum T_a_. Bats generally had lower T_sk_ when they roosted in rocks compared to in buildings or trees, *M. brandtii* had slightly colder T_sk_ during torpor bouts compared to *E. nilssonii*, and decreasing minimum T_a_ led to decreased T_sk_ (Table 3b & Fig. 3b).

## Discussion

The pre-hibernation period illustrates an important phase in the annual cycle of boreal bats, representing a critical juncture wherein they accumulate energy reserves and prepare for the impending winter. Our findings suggest that torpor use and roost selection are highly important strategies during the dynamic transition from summer to winter when faced with decreasing food availability. During the study period, *E. nilssonii* and *M. brandtii* exhibited differential roosting preferences, with *E. nilssonii* initially favoring tree roosts and later transitioning to rock screes, while *M. brandtii* shifted from favouring tree roosts to predominantly utilizing buildings.

In the late summer period (before 29th of August), torpor use was primarily predicted by the time of day, whereas later in the season (29th of August onwards), date, precipitation, and wind gained prominence in predicting torpor use. Furthermore, while the type of roost did not exert a discernible effect on the duration of torpor bouts for the species under investigation, it did impact the minimum T_sk_, with the lowest values recorded in boulder fields. Additionally, distinctive species-specific traits were observed in torpor usage and roost selection. *Eptesicus nilssonii* exhibited greater variability in roost selection and longer torpor bouts.

During the late summer, bats predominantly engaged in torpor during the daytime. This pattern aligns with the non-breeding period, where bats are exempt from expending energy on lactation or nursing pups, making energy conservation for winter hibernation a more advantageous strategy (Boyles et al., 2020). Simultaneously, this period represents a crucial phase for building energy reserves, rendering it beneficial for bats to remain active and engage in nocturnal feeding, contingent on favorable weather conditions (Fjelldal et al., 2021) and the availability of insect prey. Our late summer period dataset was constrained by the limited presence of bats on the island, as their activity typically peak around late August (Schneider & Fritzén 2020).

However, by the end of August, bats switched to using more torpor during daytime and increased torpor use in response to increasing date, higher wind speed and more rainfall. In previous studies, wind speed and rainfall have been identified to mediate state-dependent torpor responses, where individuals with higher body mass were found to increase nightly torpor use in response to poorer weather conditions, while individuals with lower body mass still used the opportunity to forage (Fjelldal et al., 2021). Flying in the rain is not only more energetically costly to the bat (Voigt et al., 2011), but also has a negative influence on food availability (Poulsen, 1996). Intriguingly, in our study, date had a stronger prediction value for torpor use than T_a_, although T_a_ has been found to be one of the most important factors in previous studies (Fjelldal et al., 2021). This suggests there are most likely other factors influencing torpor use that were not accounted for in the present study, such as food availability or function of the circannual clock.

The identified shift in torpor use in our data that occurs between mid-August and the beginning of September potentially marks the timing of when bats initiate their most rapid fat reserve accumulation, which in earlier studies has been recorded during the transition from late summer to early autumn (Kunz et al., 1998; Kronfeld-Shcor et al., 2000). It has been suggested that while temperatures and insect abundances are still elevated, insectivorous bats combine increased foraging rates with strategic torpor use in order to rapidly build sufficient energy levels for swarming and winter hibernation (Speakman et al., 1999; Fraser & McGuire, 2023). However, here we present the first results to our knowledge on how bats express torpor across this critical transition period, likely to facilitate the building of fat deposits. By increasing torpor use during daytime while still being active at night, bats should be able to maximize their energy budgets by reducing energetic expenses while maintaining a high energy intake. The period of full daytime torpor while maintaining nocturnal activity was short, and we observed that the studied bats relatively soon also begun to increase torpor use at night with the progressing autumn. The observed shift in torpor use may be in response to reaching higher fat reserve levels and thereby reducing activity levels. Unfortunately, we do not have temporal body mass data from these bats to confirm our hypothesis; however, the timing of the observed shift in torpor strategies from late summer to autumn coincides with the timing for mass gain in small bats inhabiting similar seasonal environments (Kunz et al., 1998; Kronfeld-Shcor et al., 2000).

Distinct species-specific traits were evident; notably, torpor bouts in *M. brandtii* were of shorter duration compared to bouts in *E. nilssonii*; however, this could be partly due to less data being collected on *M. brandtii* late in the season. Nevertheless, adverse weather conditions, marked by decreasing barometric pressure and T_a_, resulted in the extension of torpor bout durations in both species. The type of roost did not exert a discernible effect on the duration of torpor bouts; however, it did impact the minimum T_sk_. *Eptesicus nilssonii* has evolved to hibernate in cold climates (Solomoniv et al., 2010; Ilyukha et al., 2015), whereas *M. brandtii* demonstrates a preference for higher temperatures (Masing & Lutsar, 2007). Furthermore, the metabolic rate during torpor can exhibit variability not only between species but also within a species based on geographical origin. Individuals from northern populations may maintain a lower torpor metabolic rate at cooler temperatures compared to their southern counterparts (Dunbar & Brigham, 2010). *Myotis brandtii*, at these latitudes, is approaching its northernmost distribution limits (Tidenberg et al., 2019), potentially leading to divergent hibernating preferences compared to southern populations. Additionally, *E. nilssonii* has been observed adapting its metabolism to both high and low ambient T_a_ (Sørås et al., 2023), suggesting a potential for greater variability in thermoregulation.

*Eptesicus nilssonii* displayed greater variability in roost site selection and was observed across all three roost types, while *M. brandtii* exclusively favored buildings and tree crevices. This behavioral disparity may signify species-specific ecological adaptations, with the more southerly distributed species relying more extensively on normothermy and shelter than its counterpart accustomed to variable weather patterns throughout the active season. Notably, buildings and tree crevices serve as recognized summer roosting sites for boreal bats (Rydell, 1989; Marnell & Pretznick; Suominen et al., 2023), whereas boulder fields lack documented use during this period. Nonetheless, certain records indicate *E. nilssonii* occupying similar habitats in early spring, late autumn, and winter (Michaelsen et al., 2013; Fritzén & Hägg, 2020; Blomberg et al., 2021), suggesting these areas could serve as suitable overwintering sites. The utilization of both summer roost types and potential overwintering sites implies potential T_a_ variations in these roosts. While our study did not measure T_a_ within the roosts, it is plausible that buildings and tree crevices provide relatively warmer conditions during daytime when exposed to sun, compared to boulder fields, which are likely cooler but exhibit greater thermal stability. Additionally, the observed lower T_sk_ in bats roosting in boulder fields suggests a lower T_a_ in these locations.

The dynamic process of preparation for winter in bats involves transitioning to colder roosts and employing torpor as an energy-conserving mechanism (Speakman & Rowland, 1999). Here we show that a pivotal turning point in torpor use occurs in late summer, leading to increased reliance on daily torpor as bats likely compensate for declining insect availability during the fat-building phase. The underlying factors contributing to this turning point remain unknown and may be associated with temperature variations, rapid autumnal temperature declines at these latitudes favoring torpor in cold roosts (Speakman & Rowland, 1999), body mass considerations (Sørås et al., 2022), night length (Turner & Geiser, 2016), or a combination thereof. Moreover, after bats fully shift to employing daily torpor in early September, they also begin to increase torpor during the night. This behaviour correlates with overall decreasing T_a_ and is possibly influenced by the reduced availability of food resources in combination with individuals approaching sufficient fat reserve levels as autumn progresses (Welti et al., 2022; Speakman et al., 2000; Kronfeld-Shcor et al., 2000). Even if warm nights may occur later in autumn, insect abundance is perhaps not as substantial as at equivalent temperatures earlier in the season due to the overall diminished availability of insects. Bats during the peak fat deposit period therefore appear to respond not only to current weather conditions and food abundance, but to their own expectations of decreasing future foraging conditions. This is in accordance with the study on *M. daubentonii* by Kokurewicz & Speakman (2006), which showed that adult bats in outdoor flight-cages quickly gained body mass during the pre-hibernation fattening but then started gradually losing it again despite food being available *ad libitum*. This suggests that internal circannual clocks, temperature cues and night length may override feeding behaviour even though food could be available.

## Acknowledgments

We are grateful for Torgny Backman, Jarmo Markkanen, Katarina Meramo, Risto Lindstedt, Eeva-Maria Tidenberg, Hanna Tuominen, Ville Vasko and Johanna Yliportimo, for their hard work and help in the field catching bats. We also thank Ralf Hurst & Philipp Mallot for their assistance with raw data and Ville Vasko for his dedication on resolving Excel-problems that arise during data curation. This project was funded by Kone Foundation (grant number 201801142), Emil Aaltonen Foundation (Grant number 210219), Societas Pro Fauna Et Flora Fennica, Societas Biologica Fennica Vanamo, Ostrobotnia Australis and Waldemar von Frenckells stiftelse. Ostrobotnia Australis and Valsörarna Biological station kindly provided accommodation and transports. The radio receiver network was founded by Svensk-Österbottniska samfundet, Nordenskiöld-samfundet i Finland, Centre for Economic Development, Transport and the Environment of South Ostrobothnia and Waldemar von Frenckells stiftelse.

## Supplementary Materials 1 – breakpoint analysis

After visualizing the data, we thought there were indications of shifts in torpor use between mid-August and beginning of September (Fig. S1) and decided to perform a Davies’ test to detect potential breakpoints. When this came back significant, we used a breakpoint analysis to find an estimated breakpoint, which was detected on the day 28 after August 1^st^ (Fig. S2).

**Figure S1:**
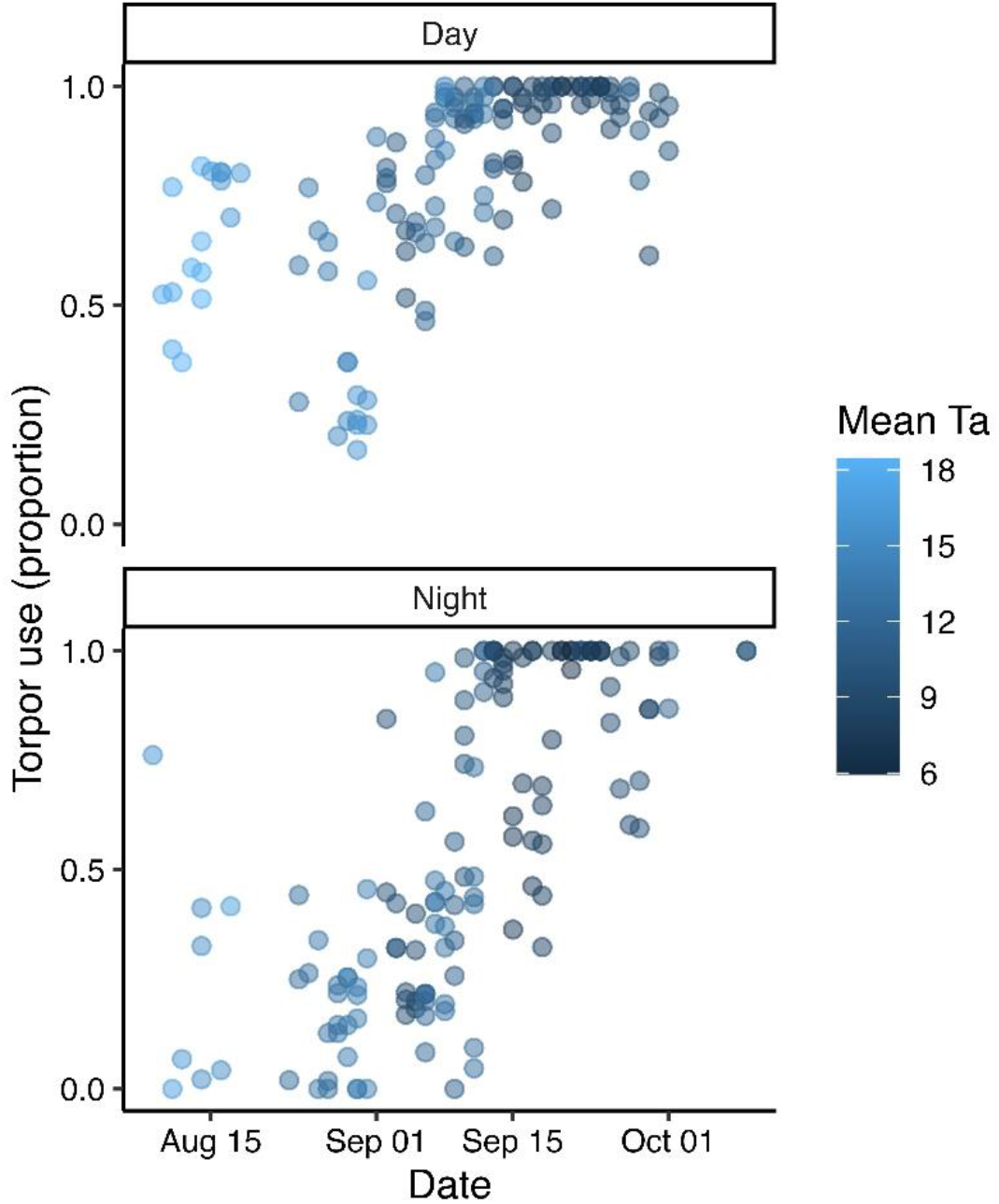
Torpor proportion data across the study period, with the color shade indicating the mean temperature during the day or night (darker blue = colder mean temperatures).

**Figure S2:**
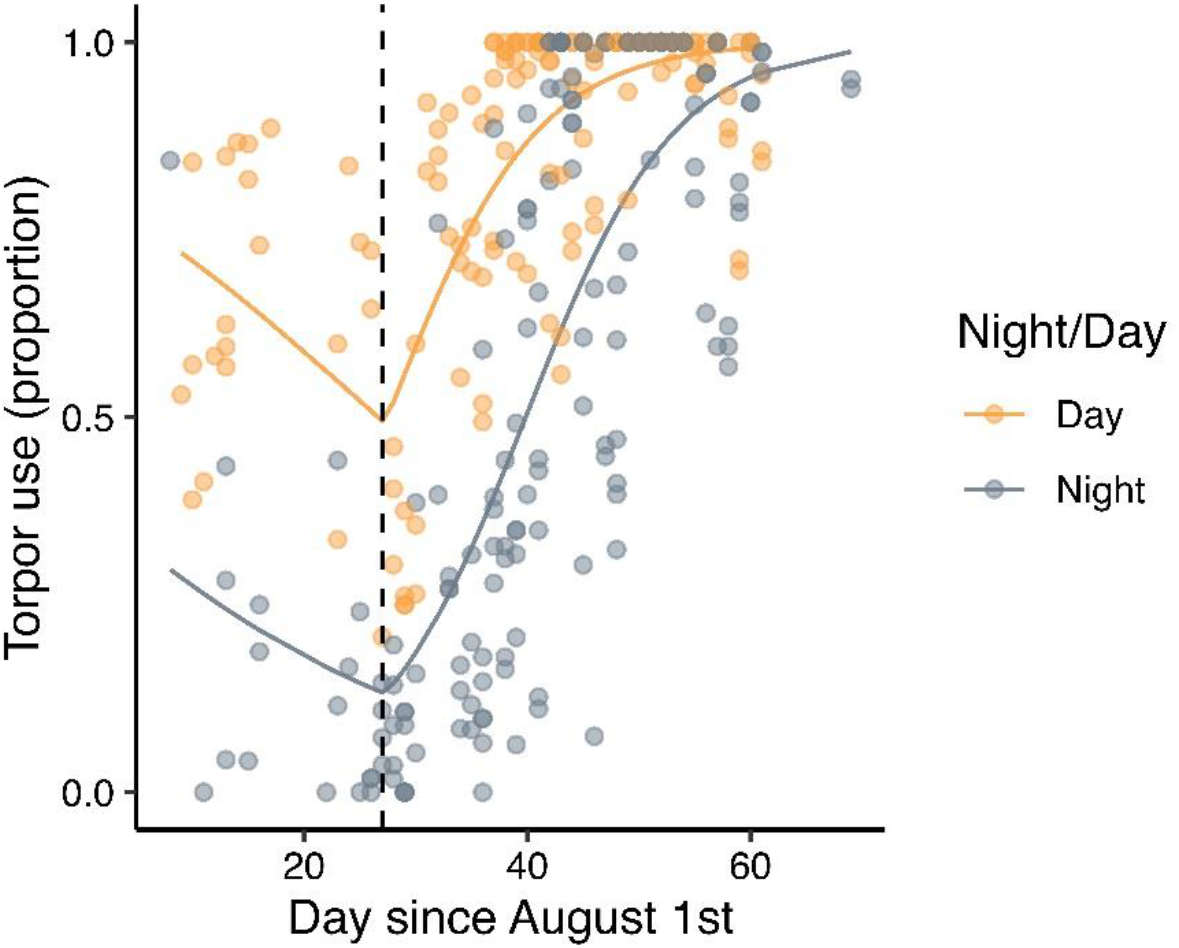
Breakpoint estimate for the torpor proportion data across the study period for nights and days. Colored lines are estimated from the breakpoint analysis model and shows the shift in torpor use at day 28 (marked with dashed line).

## Supplementary Materials 2 – model selection

**Table S1:** The 10 highest ranked models for explaining torpor use in the late summer period.

| Model | df | AICc |
| --- | --- | --- |
| *Torpor* ~ day/night | 3 | 35.9 |
| *Torpor* ~ day/night + date + species | 5 | 36.0 |
| *Torpor* ~ day/night + date | 4 | 36.5 |
| *Torpor* ~ day/night + species | 4 | 37.1 |
| *Torpor* ~ day/night + min Ta + roost + species | 6 | 37.3 |
| *Torpor* ~ day/night + roost | 4 | 37.3 |
| *Torpor* ~ day/night + min Ta | 4 | 37.4 |
| *Torpor* ~ day/night + <u>roost</u> + species | 5 | 37.5 |
| *Torpor* ~ day/night + mean_Ta | 4 | 37.8 |
| *Torpor* ~ day/night + mean Ta + species | 5 | 37.8 |

**Table S2:** The 10 highest ranked models for explaining torpor use in the autumn period.

| Model | df | AICc |
| --- | --- | --- |
| *Torpor* ~ day/night + date + wind + rain | 6 | 94.3 |
| *Torpor* ~ day/night + date + wind + rain*species | 8 | 95.1 |
| *Torpor* ~ day/night + date + wind + rain + species | 7 | 95.6 |
| *Torpor* ~ day/night + date + wind + rain + sex | 7 | 96.0 |
| *Torpor* ~ day/night + date + wind + rain + sex + species | 8 | 96.5 |
| *Torpor* ~ day/night + date + wind + rain*species + sex | 9 | 96.8 |
| *Torpor* ~ day/night + date + wind + rain + roost | 9 | 98.3 |
| *Torpor* ~ day/night + date + wind + rain + roost + sex | 10 | 99.1 |
| *Torpor* ~ day/night + date + wind + rain*species + roost | 11 | 99.5 |
| *Torpor* ~ day/night + date + wind + rain*species + roost + sex | 12 | 99.5 |

**Table S3:** The 10 highest ranked models for explaining torpor bout duration.

| Model | df | AICc |
| --- | --- | --- |
| *Duration* ~ species + min Ta + mean hPa | 6 | 1227.4 |
| *Duration* ~ species + min Ta + mean hPa + sex | 7 | 1228.5 |
| *Duration* ~ min Ta + mean hPa*species | 7 | 1229.6 |
| *Duration* ~ min Ta + sex + mean hPa*species | 8 | 1230.7 |
| *Duration* ~ min Ta + species + mean hPa*sex | 8 | 1230.7 |
| *Duration* ~ species + min Ta + mean hPa + roost | 8 | 1231.5 |
| *Duration* ~ min Ta + mean hPa + species*roost | 9 | 1231.7 |
| *Duration* ~ species + min Ta + mean hPa + sex + roost | 9 | 1232.2 |
| *Duration* ~ min Ta + mean hPa | 5 | 1232.4 |
| *Duration* ~ min Ta + mean hPa + sex + species*roost | 10 | 1232.4 |

**Table S4:**
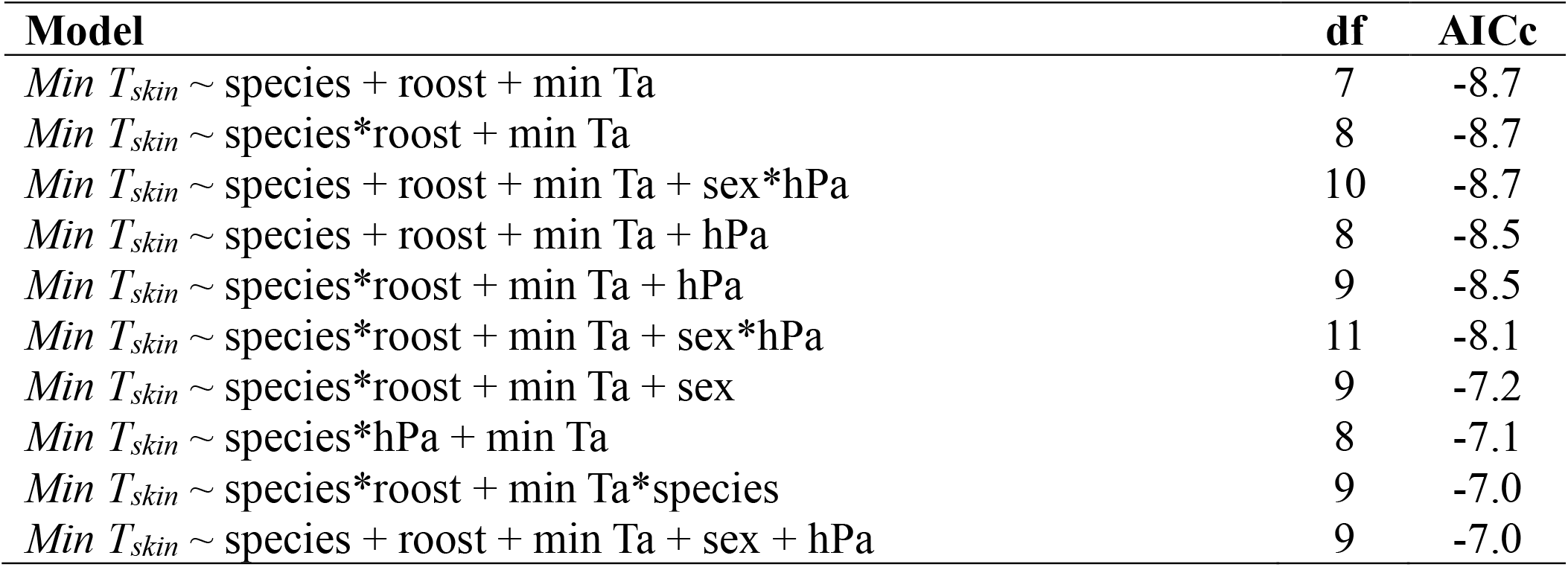
The 10 highest ranked models for explaining minimum T_sk_ during torpor.

## References

Aldridge, H. D. J. N., & Brigham, R. M. (1988). Load-carrying and maneuverability in an insectivorous bat: a test of the 5% ‘rule’ of radiotelemetry. Journal of Mammalogy, 69(2), 379–382.

Anthony, E. L. P. (1988). Age determination in bats. In Ecological and behavioral methods for the study of bats:31–46. Kunz, T. (Ed.). Washington, DC: Smithsonian Institute Press.

Audet, D., & Thomas, D. W. (1996). Evaluation of the accuracy of body temperature measurement using external radio transmitters. Canadian Journal of Zoology, 74, 1778–1781.

Barclay, R. M., Lausen, C. L., & Hollis, L. (2001). What’s hot and what’s not: defining torpor in free-ranging birds and mammals. Canadian Journal of Zoology, 79(10), 1885–1890.

Barton, K., & Barton, M. K. (2015). Package ‘mumin’. Version, 1, 439.

Bates, D., Mächler, M., Bolker, B., & Walker, S. (2015). Fitting linear mixed-effects models using lme4. Journal of Statistical Software, 67, 1–48.

Blomberg, A. S., Vasko, V., Meierhofer, M. B., Johnson, J. S., Eeva, T., & Lilley, T. M. (2021). Winter activity of boreal bats. Mammalian Biology, 101, 609–618.

Bondarenco, A., Körtner, G., & Geiser, F. (2016). How to keep cool in a hot desert: torpor in two species of free-ranging bats in summer. Temperature, 3(3), 476–483.

Boyles, J. G. (2017). Benefits of knowing the costs of disturbance to hibernating bats. Wildlife Society Bulletin, 41(2), 388–392.

Boyles, J. G., Johnson, J. S., Blomberg, A., & Lilley, T. M. (2020). Optimal hibernation theory. Mammal Review, 50, 91–100

Czenze, Z. J., Smit, B., van Jaarsveld, B., Freeman, M. T., & McKechnie, A. E. (2022). Caves, crevices and cooling capacity: Roost microclimate predicts heat tolerance in bats. Functional Ecology, 36, 38–50. 10.1111/1365-2435.13918

Czenze, Z. J., Jonasson, K. A., & Willis, C. K. (2017). Thrifty females, frisky males: winter energetics of hibernating bats from a cold climate. Physiological and Biochemical Zoology, 90(4), 502–511.

Dunbar, M. B., & Brigham, R. M. (2010). Thermoregulatory variation among populations of bats along a latitudinal gradient. Journal of Comparative Physiology B, 180, 885–893. 10.1007/s00360-010-0457-y

Fjelldal, M. A., Wright, J., & Stawski, C. (2021). Nightly torpor use in response to weather conditions and individual state in an insectivorous bat. Oecologia, 197(1), 129–142.

Fjelldal, M. A., Sørås, R., & Stawski, C. (2022). Universality of torpor expression in bats. Physiological and Biochemical Zoology, 95(4), 326–339.

Fjelldal, M. A., Stawski, C., Sørås, R., & Wright, J. (2023a). Determining the different phases of torpor from skin-or body temperature data in heterotherms. Journal of Thermal Biology, 111, 103396. 10.1016/j.jtherbio.2022.103396

Fjelldal, M. A., Muller, A. S., Ratikainen, I. I., Stawski, C., & Wright, J. (2023b). The small-bat-in-summer paradigm: Energetics and adaptive behavioural routines of bats investigated through a stochastic dynamic model. Journal of Animal Ecology, 92(10), 2078–2093.

Fraser, E. E., & McGuire, L. P. (2023). Prehibernation swarming in temperate bats: a critical transition between summer activity and hibernation. Canadian Journal of Zoology.

Fritzén, N., & Hägg, J. (2020). Valsörarnas biologiska station – verksamhetsberättelse för år 2019. OA-Natur, 22, 3–17.

Geiser, F. (1988). Reduction of metabolism during hibernation and daily torpor in mammals and birds: temperature effect or physiological inhibition? Journal of Comparative Physiology B, 158(1), 25–37.

Geiser, F., & Ruf, T. (1995). Hibernation versus daily torpor in mammals and birds: physiological variables and classification of torpor patterns. Physiological zoology, 68(6), 935–966.

Geiser, F., & Brigham, R. (2000). Torpor, thermal biology, and energetics in Australian long-eared bats (Nyctophilus). Journal of Comparative Physiology B, 170, 153–162. 10.1007/s003600050270

Geiser, F. (2004). Metabolic rate and body temperature reduction during hibernation and daily torpor. Annual Review of Physiology, 66, 239–274. 10.1146/annurev.physiol.66.032102.115105

Geiser, F., Currie, S. E., O’Shea, K. A., & Hiebert, S. M. (2014). Torpor and hypothermia: reversed hysteresis of metabolic rate and body temperature. American Journal of Physiology-Regulatory, Integrative and Comparative Physiology, 307, R1324–R1329.

Gottwald, J., Ziedler, R., Friess, N., Marvin, L., Rudenbach, C., & Nauss, T. (2019). Introduction of an automatic and open-source radio-tracking system for small animals. Methods in Ecology and Evolution, 00(0), 1–10. DOI: 10.1111/2041-210X.13294

Humphries, M. M., Kramer, D. L., & Thomas, D. W. (2003). The role of energy availability in Mammalian hibernation: an experimental test in free-ranging eastern chipmunks. Physiological and Biochemical Zoology, 76(2), 180–186. 10.1086/367949

Ilyukha, V. A., Antonova, E., Belkin, V. A., Uzenbaeva, L., Khizhkin, E., Sergina, S., Ilyina, T., Baishnikova, I., Kizhina, A. (2015). The eco-physiological status of hibernating bats (Chiroptera) in the north of the European distribution range. Acta Biologica Universitatis Daugavpiliensis, 15, 75–94.

Kokurewicz, T., & Speakman, J. R. (2006). Age related variation in the energy costs of torpor in Daubenton’s bat: effects on fat accumulation prior to hibernation. Acta Chiropterologica, 8, 509–521.

Kotila, M., Suominen, K. M., Vasko, V. V., Blomberg, A. S., Lehikoinen, A., Andersson, T., … Lilley, T. M. (2023). Large-scale long-term passive-acoustic monitoring reveals spatio-temporal activity patterns of boreal bats. Ecography, 2023, e06617. 10.1111/ecog.06617

Kronfeld-Schor, N., Richardson, C., Silvia, B. A., Kunz, T. H., & Widmaier, E. P. (2000). Dissociation of leptin secretion and adiposity during prehibernatory fattening in little brown bats. American Journal of Physiology-Regulatory, Integrative and Comparative Physiology, 279, R1277–R1281.

Kunz, T. H., Wrazen, J. A., & Burnett, C. D. (1998). Changes in body mass and fat reserves in pre-hibernating little brown bats (Myotis lucifugus). Ecoscience, 5, 8–17.

Marnell, F., & P. Presetnik, P. (2010). Protection of overground roosts for bats (particularly roosts in buildings of cultural heritage importance). EUROBATS Publication Series No. 4 (English version). UNEP/EUROBATS Secretariat, Bonn, Germany (2010), p. 57.

Masing, M., & Lutsar, L. (2007). Hibernation temperatures in seven species of sedentary bats (Chiroptera) in north-eastern Europe. Acta Zool Litu, 17, 47–55. 10.1080/13921657.2007.10512815

Michaelsen, T., Olsen, O., & Grimstad, K. (2013). Roosts used by bats in late autumn and winter at northern latitudes in Norway. Folia Zool, 62, 297–303. 10.173325225/fozo.v62.i4.a7.2013

Mitchell-Jones, T., & McLeish, A. (2003). Bat Worker’s Manual (3rd ed.). Joint Nature Conservation Committee, p. 178.

Muggeo, V. M. (2008). Segmented: an R package to fit regression models with broken-line relationships. R news, 8, 20–25.

Muñoz-Garcia, A., Ben-Hamo, M., Pilosof, S., Williams, J. B., & Korine, C. (2022). Habitat aridity as a determinant of the trade-off between water conservation and evaporative heat loss in bats. Journal of Comparative Physiology B, 192(2), 325–333.

Poulsen BO (1996). Relationships between frequency of mixed-species flocks, weather and insect activity in a montane cloud forest in Ecuador. Ibis, 138, 466–470.

Rubalcaba, J. G., Gouveia, S. F., Villalobos, F., Cruz-Neto, A. P., Castro, M. G., Amado, T. F., … & Olalla-Tárraga, M. Á. (2022). Physical constraints on thermoregulation and flight drive morphological evolution in bats. Proceedings of the National Academy of Sciences, 119(15), e2103745119.

Ruf, T., & Geiser, F. (2015). Daily torpor and hibernation in birds and mammals. Biological Reviews, 90(3), 891–926.

Rydell, J. (1989). Feeding activity of the northern bat Eptesicus nilssoni during pregnancy and lactation. Oecologia, 80, 562–565. 10.1007/BF00380082

Schneider, M., & Fritzén, N. R. (2020). Flador och deras insektproduktion – betydelsen för lokala och migrerande fladdermöss i Kvarken. - Delrapport inom Interreg Botnia Atlantica projekt Kvarken Flada. 72 s.

Solomonov NG, Anufriev AI, Solomonova TN (2010) Mechanisms of hibernation in small mammals of Yakutia. Cryobiology, 61, 397. 10.1016/j.cryobiol.2010.10.120

Speakman J. R. & Rowland A. (1999). Preparing for inactivity: how insectivorous bats deposit a fat store for hibernation. Proc. Nutr. Soc., 58, 123–131.

Speakman, J. R., Rydell, J., Webb, P. I., Hayes, J. P., Hays, G. C., Hulbert, I. A. R., and Mcdevit, R. M. (2000). Activity patterns of insectivorous bats and birds in northern Scandinavia (69°N), during continuous midsummer daylight. Oikos, 88, 75–86. 10.1034/j.1600-0706.2000.880109.x

Speakman, J. R., & Thomas, D. W. (2003). Physiological ecology and energetics of bats in Kunz, T. H., & Fenton, M. B. (2003): Bat ecology, pp. 430–490.

Stawski C, Geiser F (2010) Seasonality of torpor patterns and physiological variables of a free-ranging subtropical bat. J Exp Biol, 213, 393–399. 10.1242/jeb.038224

Suominen, K. M., Kotila, M., Blomberg, A. S., Pihlström, H., Ilyukha, V., & Lilley, T. M. (2022). Northern bat Eptesicus nilssonii (Keyserling and Blasius, 1839). In Handbook of the mammals of Europe (pp. 1–27). Cham: Springer International Publishing.

Suominen, K. M., Vesterinen, E. J., Kivistö, I., Reiman, M., Virtanen, T., Meierhofer, M. B., … & Lilley, T. M. (2023). Environmental features around roost sites drive species-specific roost preferences for boreal bats. Global Ecology and Conservation, e02589.

Sørås, R., Fjelldal, M. A., Bech, C., van der Kooij, J., Skåra, K. H., Eldegard, K., & Stawski, C. (2022). State dependence of arousal from torpor in brown long-eared bats (Plecotus auritus). Journal of Comparative Physiology B, 192(6), 815–827.

Sørås, R., Fjelldal, M. A., Bech, C., van der Kooij, J., Eldegard, K., & Stawski, C. (2023). High latitude northern bats (Eptesicus nilssonii) reveal adaptations to both high and low ambient temperatures. Journal of Experimental Biology, 226(21).

Thieurmel, B., & Elmarhraoui, A. (2019). suncalc: Compute sun position, sunlight phases, moon position and lunar phase. R package version 0.5.0. Retrieved from https://CRAN.R-project.org/package=suncalc

Tidenberg, E.-M., Liukko, U.-M., & Stjernberg, T. (2019): Atlas of Finnish bats. Ann Zool Fenn, 56, 207–250.

Turner, J. M., & Geiser, F. (2017). The influence of natural photoperiod on seasonal torpor expression of two opportunistic marsupial hibernators. J Comp Physiol B, 187, 375–383. 10.1007/s00360-016-1031-z

Voigt, C. C., Schneeberger, K., Voigt-Heucke, S. L., & Lewanzik, D. (2011). Rain increases the energy cost of bat flight. Biology letters, 7(5), 793–795.

Welti, E. A., Zajicek, P., Frenzel, M., Ayasse, M., Bornholdt, T., Buse, J., … & Haase, P. (2022). Temperature drives variation in flying insect biomass across a German malaise trap network. Insect conservation and diversity, 15(2), 168–180.

Willis, C. K. R. (2007). An Energy-Based Body Temperature Threshold between Torpor and Normothermia for Small Mammals. Physiological and Biochemical Zoology, 80(6), 643–651. doi:10.1086/521085

